# Neural entrainment predicts the anticipatory P300 component during musical meter perception: An EEG study using dual-meter sound stimuli

**DOI:** 10.1101/2025.07.01.662512

**Authors:** Sotaro Kondoh, Kazuo Okanoya, Ryosuke O. Tachibana

## Abstract

Meter, the organization of beats into regular groupings, is a core element of music perception that arises from the interplay between the bottom-up processing of acoustic features and top-down attentional mechanisms. To disentangle this interplay, we previously developed dual-meter stimuli consisting of band-limited noise bursts that varied in center frequency and duration, with one feature following a triple-meter pattern and the other a quadruple-meter pattern. The perceived meter thus switches between the two by shifting attention from one acoustic feature to another, even when the physical stimulus remains identical. Using these stimuli, we investigated whether directing attention to specific acoustic features is associated with neural entrainment and anticipatory processing of the perceived metrical structure, and whether the former predicts the latter, by recording electroencephalogram (EEG) signals while isolating the effects of acoustic differences. We found that EEG spectral profiles shifted toward the metrical structure of the attended feature, and that the P300 component, an index of anticipatory processing, tended to be larger when the metrical structure was disrupted than when it remained intact, although both measures varied considerably across individuals. Critically, linear regression analysis revealed that participants whose neural entrainment more closely tracked the attended metrical structure exhibited significantly larger P300 responses to metrical violations. These findings suggest that attending to specific acoustic features links neural entrainment with anticipatory processing, which may underlie meter perception.

## 1. Introduction

Meter is one of the core elements of music perception. It emerges from the listener’s ability to extract regular beats and assign varying levels of subjective intensity to them, thereby forming organized groupings (Cooper & Meyer, 1963; Fujii & Schlaug, 2013; Honing, 2013; Kotz et al., 2018; Large & Jones, 1999; Lerdahl & Jackendoff, 1983; London, 2012; Palmer & Krumhansl, 1990; Patel, 2006, 2010). In a duple meter, as found in march music, beats follow a recurring “strong(s)-weak(w)-s-w-…” pattern. The triple meter organizes beats as “s-w-w-s-w-w-…,” which typically characterizes waltz music. Although the meter is a familiar aspect of musical experience, the processes by which listeners extract and maintain such metrical structures from complex auditory input remain not fully understood.

Meter perception involves both the bottom-up processing of acoustic features and top-down attentional mechanisms. Salient acoustic regularities can drive meter perception in a bottom-up manner. For instance, a sound sequence such as “loud-soft-loud-soft…” is perceived as a duple meter, with louder sounds marking strong beats (Chen et al., 2006, 2009; Snyder & Large, 2005; Zatorre et al., 2007). However, music can also contain multiple acoustic cues that support different metrical groupings at the same time. When these cues favor conflicting metrical interpretations, listeners must selectively attend to particular features in order to perceive a coherent meter. This process akin to auditory scene analysis, in which competing acoustic organizations are resolved through attentional mechanisms (Bregman, 1994; Bregman et al., 2001; Cherry, 1953). To investigate the role of top-down attention in meter perception, we previously developed “dual-meter” stimuli consisting of sequences of band-limited noise bursts with varying center frequencies and durations (Kondoh et al., 2021). In these stimuli, frequency and duration patterns simultaneously supported different meters: for example, a “high–low–low…” frequency pattern established a triple meter, whereas a “long–short–short–short…” duration pattern corresponded to a quadruple meter. When participants were instructed to shift their attention between frequency and duration, they experienced different meters even though the physical sound sequences remained unchanged. These findings support the contribution of top-down processing to meter perception.

Given that top-down attention shapes meter perception, a critical question arises regarding the neural mechanisms underlying this process. Recent research suggests that neural entrainment and anticipatory processing are key mechanisms (Harding et al., 2025; Snyder et al., 2024; Vuust et al., 2022). Neural entrainment refers to the synchronization of endogenous brain oscillations with periodic auditory input (Fujioka et al., 2009; Nozaradan et al., 2011). Although this synchronization is driven by the acoustic regularities of the stimulus, dynamic attending theory proposes that neural entrainment also generates expectations about upcoming events, providing a foundation for anticipatory processing (Ellis & Jones, 2009; Jones, 2018; Jones & Boltz, 1989). Integrating these ideas with the role of top-down attention in meter perception, we hypothesized that selectively attending to a particular acoustic feature enhances neural entrainment to the regularity defined by that feature, and that this entrainment, in turn, gives rise to anticipation of the corresponding metrical structure. However, direct evidence for this proposed link has been difficult to obtain because, in conventional paradigms, acoustic feature differences and perceived metrical accents are confounded, making it impossible to determine whether the observed neural responses reflect meter perception or the bottom-up processing of acoustic differences. The dual-meter paradigm offers a means of overcoming this limitation. Because the physical stimulus remains constant across attentional conditions, comparing neural activity when listeners attend to different acoustic features enables the isolation of meter-related entrainment and anticipation from responses driven by the acoustic input.

To examine this hypothesis, the present study combined the dual-meter paradigm with electroencephalogram (EEG) recordings. We first assessed whether neural entrainment and anticipatory processing could each be observed independently and then tested whether the strength of entrainment predicted the magnitude of anticipatory responses. For neural entrainment, we compared the spectral power of EEG when participants attended to either the frequency or duration of dual-meter stimuli. For anticipatory processing, we analyzed event-related potentials (ERPs) elicited by violations of metrical expectations introduced by altering the frequency or duration of a single noise burst in the stimuli. Such violations evoke mismatch negativity (MMN), which reflects the automatic detection of auditory deviations, and P300, which is associated with the conscious evaluation of unexpected events (Koelsch et al., 2019; Näätänen et al., 1978; Sutton et al., 1965; Vuust et al., 2009). Finally, we examined whether meter-related neural entrainment predicted ERP responses to violations of the anticipated metrical structure.

## 2. Methods

### 2.1. Participants

Thirty-four participants (17 males, 17 females; 19.8 ± 1.6 years old) were recruited for this study from March 9 to October 1, 2022. All participants reported normal hearing and no history of hearing impairment. Before participation, they received instructions regarding the study and signed a written informed consent form. This study was approved by the Ethical Review Committee for Experimental Research on Human Subjects at the Graduate School of Arts and Sciences, the University of Tokyo, Japan (#663).

The sample size was determined a priori using G*Power 3 (Faul et al., 2007). The primary basis for sample size estimation was within-subject comparisons, including pairwise comparisons of neural entrainment indices across meter-related frequency bands and ERP amplitude differences between violated and unviolated conditions (see ***2.5.4. Statistics***). Based on a pilot experiment that yielded a medium effect size (Cohen’s *d* = 0.51) for ERP amplitude differences, a two-tailed paired *t*-test with *α* = 0.05 and *power* = 0.80 indicated a required sample size of 34 participants. Additionally, we conducted a sensitivity power analysis (Lakens, 2022) for the regression analysis to examine whether neural entrainment indices predicted ERP responses to violations of metric anticipation. With 34 participants, *α* = 0.05, and *power* = 0.80, the linear regression with one predictor was powered to detect effect sizes of f^2^ = 0.25.

### 2.2. Stimuli

We designed dual-meter stimuli (**Fig. 1A, B**) consisting of 24 band-noise bursts. Each burst had one of two center frequencies: a high frequency of 1600 Hz or a low frequency of 1400 Hz, resulting in a frequency difference of 200 Hz. We also varied the duration of the bursts and classified them as long (150 ms) or short (50 ms) sounds, creating a duration difference of 100 ms. These specific frequency and duration differences were selected based on our previous study (Kondoh et al., 2021), which demonstrated that listeners were most likely to switch attention between acoustic features under these parameters. The inter-onset intervals (IOIs) were fixed at 300 ms. The bandwidth of the noise bursts was 120% of the center frequency, with rise and decay times of 10 ms.

**Fig 1.**
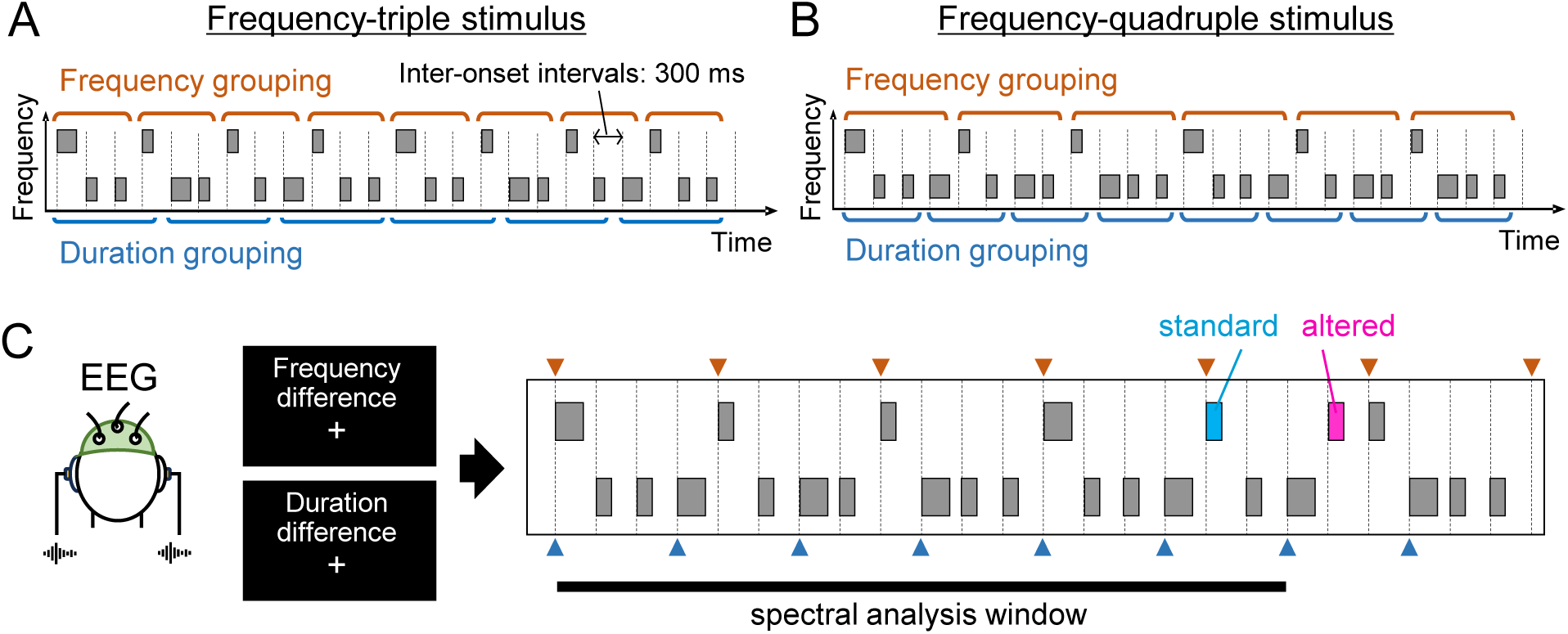
Schematic drawing of dual-meter stimuli and experimental design. **A.** Example stimulus from the frequency-triple category with “high-low-low-…” frequency and “long-short-short-short-…” duration patterns. **B.** Example stimulus from the frequency-quadruple category with “high-low-low-low…” frequency and “long-short-short-…” duration patterns. See **Supplementary Fig. S1** for all stimulus types. The center frequency of the higher sound was 1600 Hz and that of the lower sound was 1400 Hz, resulting in a frequency difference of 200 Hz. The duration of the longer sound was 150 ms, whereas that of the shorter sound was 50 ms, creating a duration difference of 100 ms. The bandwidth of each noise burst was 120% of its center frequency, and the inter-onset intervals (IOIs) were fixed at 300 ms. **C.** Experimental procedure. Participants attended to either the duration or frequency differences of the stimulus during electroencephalogram (EEG) recordings. Spectral analysis was performed on the EEG data corresponding to the initial 18 sounds (indicated by the black line). In this example, the 20th noise burst was altered by increasing its center frequency from low to high (altered sound is indicated by the pink square). We defined standard sounds as those that matched the altered sounds in center frequency and duration, presenting at least three sounds earlier (indicated by the light blue square). When focusing on the frequency difference, the metrical structure of the quadruple meter (indicated by orange triangles) is violated by the altered sound (violated condition). In contrast, when focusing on the duration difference, the metrical structure of the triple meter (indicated by blue triangles) remains intact (unviolated condition). Event-related potentials (ERPs) were calculated around the altered and standard sounds. See **Supplementary Fig. S2** for all patterns of alternation.

We created two types of stimuli based on regularity patterns in center frequency and duration: frequency-triple stimuli (**Fig. 1A**), and frequency-quadruple stimuli (**Fig. 1B**). Participants were randomly assigned to one of these two stimulus categories, with half listening to the frequency-triple stimuli and the other half listening to the frequency-quadruple stimuli. In the frequency-triple condition, the frequency pattern repeated every three sounds (“high–low–low” or “low–high–high”), while the duration pattern repeated every four sounds (“long–short–short–short” or “short–long–long–long”). In contrast, the frequency-quadruple category featured a frequency pattern that repeated every four sounds (“high–low–low–low” or “low–high–high–high”), paired with a duration pattern that repeated every three sounds (“long–short–short” or “short–long–long”). Consequently, each category contained four stimulus subtypes determined by which sounds served as strong beats: high and long, high and short, low and long, and low and short (see **Supplementary Fig. S1** for all stimulus types).

### 2.3. Pre-training to familiarize with dual-meter stimuli

Participants were familiarized with dual-meter stimuli for focusing on center frequency and duration differences by attending a web-based pre-training session at their home. In this pre-training, they were instructed to listen to a dual-meter stimulus and count specific target sounds. The target sound was instructed by showing a word, “high,” “low,” “long,” or “short,” on the computer screen before starting the stimulus presentation. The task of participants was to detect the instructed sound in the stimulus and count them. Each stimulus contained 3 to 32 sounds per trial, and included one to eight target sounds. After listening, the participants reported the target count by selecting numbers from 1 to 8 on the screen.

The training consisted of six blocks of 32 trials (4 characteristics × 8 targets). In the high/low counting trials, the frequency difference remained at 200 Hz, whereas the duration difference increased from 64 ms in the first and second blocks to 80 ms in the third and fourth blocks, rendering the frequency difference less prominent. Conversely, in the long/short counting trials, the duration difference remained at 100 ms, whereas the frequency difference increased from 128 Hz in the first and second blocks to 160 Hz in the third and fourth blocks, making the duration difference less pronounced. In the final two blocks, both tasks used stimuli with a frequency difference of 200 Hz and a duration difference of 100 ms, matching the parameters used for EEG recording. No feedback on correct answers was provided during training. The training program was developed using lab.js (Henninger et al., 2022), and the participants completed the program individually on their personal computers with headphones. To verify the use of headphones, we used a pre-set screening test (Woods et al., 2017).

### 2.4. EEG recording

During the EEG recording session, the participants were asked to listen to the dual-meter stimuli while attending to differences in either frequency or duration without any explicit response (**Fig. 1C**). To violate metrical anticipation, the center frequency or duration of the 19th, 20th, or 21st band-noise bursts was modified (‘altered’ in **Fig. 1C**; see **Supplementary Fig. S2** for all alterations). For comparison, ‘standard’ sounds were defined as those with both center frequency and duration matching the altered sounds. We further discriminated the following two situations: one was the ‘violated’ condition, where changes in the attended acoustic feature disrupted metrical anticipation, and the other was the ‘unviolated’ condition, where the metrical structure remained intact despite modifications.

Each trial began with 0.8 seconds of silence followed by a 7.2-second stimulus presentation. A block consisted of 14 trials that focused on one acoustic feature, including six trials for each of the violated and unviolated conditions, and two trials for stimuli with no alteration. Throughout each block, a fixation cross and a display indicating the target feature were shown on the screen. Each session comprised four blocks with randomized stimulus assignments and trial orders, and the participants completed a total of four sessions.

EEG recordings were performed using a 32-channel EasyCap and BrainAmp DC system (Brain Products GmbH, Germany) at a sampling rate of 1000 Hz. The electrodes were positioned at C3, C4, CP1, CP2, CP5, CP6, Cz, F3, F4, F7, F8, FC1, FC2, FC5, FC6, FT10, Fp1, Fp2, Fz, Iz, O1, O2, P3, P4, P7, P8, PO10, PO9, Pz, T7, and T8. In addition, an electrode (AFp10) was placed near the right eye to record electrooculogram activity, and reference electrodes were attached to the left (A1) and right (A2) earlobes. The stimulus presentation was controlled using Presentation 18.1 software (Neurobehavioral Systems, Inc., USA). Instructions regarding the attended acoustic feature and the fixation cross were presented on a 24-inch LCD monitor (XL2420T, BenQ, Taiwan), with participants seated approximately 80 cm from the screen. Auditory stimuli were delivered binaurally through earphones (Compumedics Neuroscan Inc., USA) at approximately 70 dB sound pressure level (SPL) via an external sound card (UA-22, Roland, Japan).

### 2.5. Analysis

#### 2.5.1. Preprocessing on EEG signal

Preprocessing was performed using MATLAB R2023b and EEGLAB v2022.1 (Delorme & Makeig, 2004). EEG data obtained from electrodes with impedance values greater than 15 kΩ were excluded from the analysis. The raw signal was filtered using a high-pass filter set at 0.5 Hz and a low-pass filter set at 30 Hz. Independent component analysis (ICA) was then performed to decompose the EEG data into a number of components equal to the number of electrodes (Delorme et al., 2007; Makeig et al., 2004). The ICLabel tool was used to classify these components as either brain-derived signals or noise sources, including ocular, muscular, and cardiac activities and line noise (Pion-Tonachini et al., 2019). For subsequent analyses, only components exhibiting more than 50% brain-derived signals were retained.

#### 2.5.2. Neural entrainment to meter

The neural entrainment to meter patterns was assessed by the following three steps: obtaining spectra, selecting appropriate electrodes, and calculating meter-related components. EEG data corresponding to the first 18 noise bursts of the dual-meter stimuli (total 5.4 s) in each trial were used to assess neural entrainment. A moving median filter with a 1.2-second window was applied to detect and eliminate slow drifts. The beginning and last parts of the EEG signal were then tapered using a raised cosine (rise time: 0.9 s; decay time: 0.3 s) to reduce side-lobe effects in the spectra. The data were downsampled to a sampling rate of 40 Hz for computational efficiency. Then, a Fast Fourier Transform (FFT) was performed at 1,728 points (with 8-time interpolation) to obtain amplitude spectra with a frequency resolution at 0.023 Hz (40 Hz / 1,728 points).

We then selected appropriate electrodes by identifying neural oscillations with a frequency of the stimulus IOI (300 ms). We compared the grand means of the EEG spectra with the surrogate baseline spectrum for each electrode. This surrogate signal was generated by randomly shuffling the original EEG data for every 300-ms time slots, and underwent the same tapering and FFT processes (**Fig. 2A**). For each electrode, we computed the amplitude ratio of the spectral component at 3.33 Hz (ranging from 2.83 to 3.83 Hz) in the original signal to that in the surrogate signal. We selected an electrode with the highest ratio and surrounding electrodes and averaged their signals for further neural entrainment analysis (**Fig. 2B**).

**Fig. 2.**
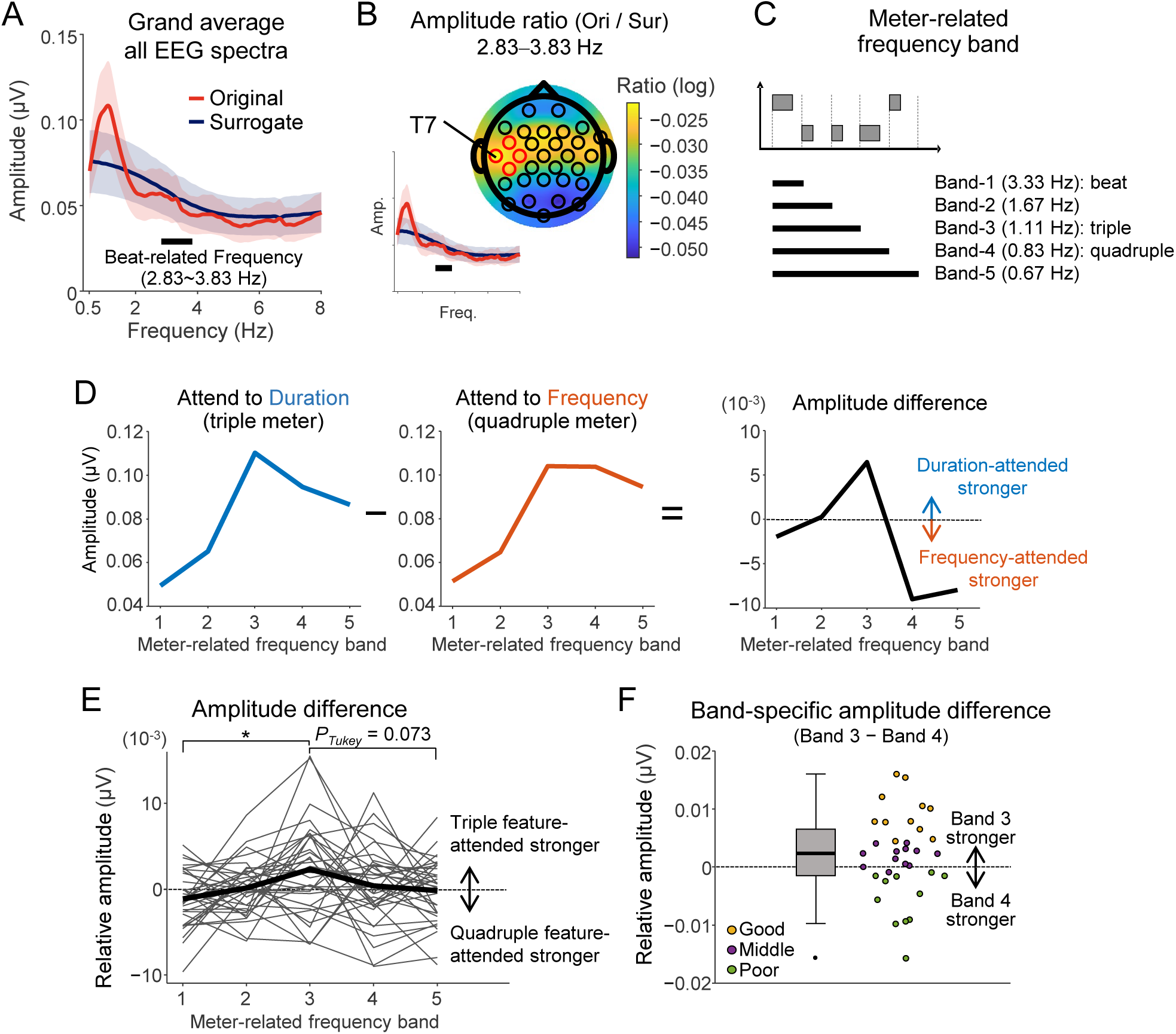
Neural index reflecting entrainment associated with meter perception. **A.** Grand average of EEG spectra of the original and surrogate signals without differentiating whether participants focused on triple- or quadruple-related acoustic features. The shaded areas represent the standard deviations (SD) across the electrodes. The analysis targeted a beat-related frequency range (2.83–3.83 Hz). **B.** Topographical electrode distribution of the log-transformed ratio of the original to surrogate EEG signal amplitudes in the beat-related frequency range. The T7 electrode showed the highest ratio, and the adjacent electrodes, FC5, C3, and CP5, were selected for further analysis (indicated by red circles). **C.** Description of meter-related frequency bands: Band-1 (3.33 [3.26–3.43] Hz) corresponding to beat, Band-2 (1.67 [1.60–1.76] Hz), Band-3 (1.11 [1.04–1.20] Hz) corresponding to the triple meter, Band-4 (0.83 [0.76–0.93] Hz) corresponding to the quadruple meter, and Band-5 (0.67 [0.60–0.76] Hz). **D.** Spectral analysis to calculate neural entrainment (examples from one participant). We calculated the spectral amplitude for the triple-related (left) and quadruple-related features (middle) and computed the difference (triple minus quadruple) to eliminate the effects of acoustic variability (right). **E.** Overall amplitude differences between triple-related and quadruple-related acoustic features across meter-related frequency bands. Each grey line represents individual data, whereas the bold black line indicates the average across participants. Positive values indicate stronger entrainment to triple-related features, whereas negative values indicate stronger entrainment to quadruple-related features. Asterisk (*) indicates a significant difference (*p_Tukey_* < 0.05). **F.** Box plots and individual data of the band-specific amplitude difference between Bands 3 and 4. Greater values indicate stronger entrainment to triple-related features in Band-3 and weaker entrainment to quadruple-related features in the same band. The participants were classified into three groups in descending order: good (*n* = 11), middle (*n* = 12), and poor switchers (*n* = 11) for visualization purpose (see Fig. 3F**-H**).

After selecting the electrodes, we calculated an index reflecting meter-related neural entrainment. We defined five frequency bands (Band-1 to Band-5), centered at 3.33 Hz (range 3.26–3.43 Hz), 1.67 Hz (1.60–1.76 Hz), 1.11 Hz (1.04–1.20 Hz), 0.83 Hz (0.76–0.93 Hz), and 0.67 Hz (0.60–0.76 Hz), respectively (**Fig. 2C**). Band-1, 3, and 4 correspond to the regularities of the beat, triple meter, and quadruple meter, respectively. We computed the amplitude of each band in the mean EEG signal from the selected electrodes when participants attended to triple-related or quadruple-related features and then calculated the difference between them (**Fig. 2D**). The amplitude difference is more positive when there is stronger neural entrainment to the triple feature. It is more negative when there is stronger entrainment to the quadruple feature.

#### 2.5.3. Event-related potentials

For the analysis of event-related potentials associated with metrical anticipation, we selected appropriate electrodes by identifying strong neural responses to the altered sound in stimulus sequence. We computed the average signals for each of standard and altered sounds irrespective of the difference between violated and unviolated conditions (**Fig. 3A**), applying the analysis window of 0–600 ms after the sound onset with the baseline correction using the mean amplitude between 100 and 0 ms prior to sound onset. The electrode exhibiting the largest difference between the standard and altered sounds, along with its neighboring electrodes, was chosen for further ERP analysis (**Fig. 3B**). We employed this selection criterion to ensure that the EEG data contained the neural response that reflected the bottom-up process for detecting acoustic differences.

**Fig. 3.**
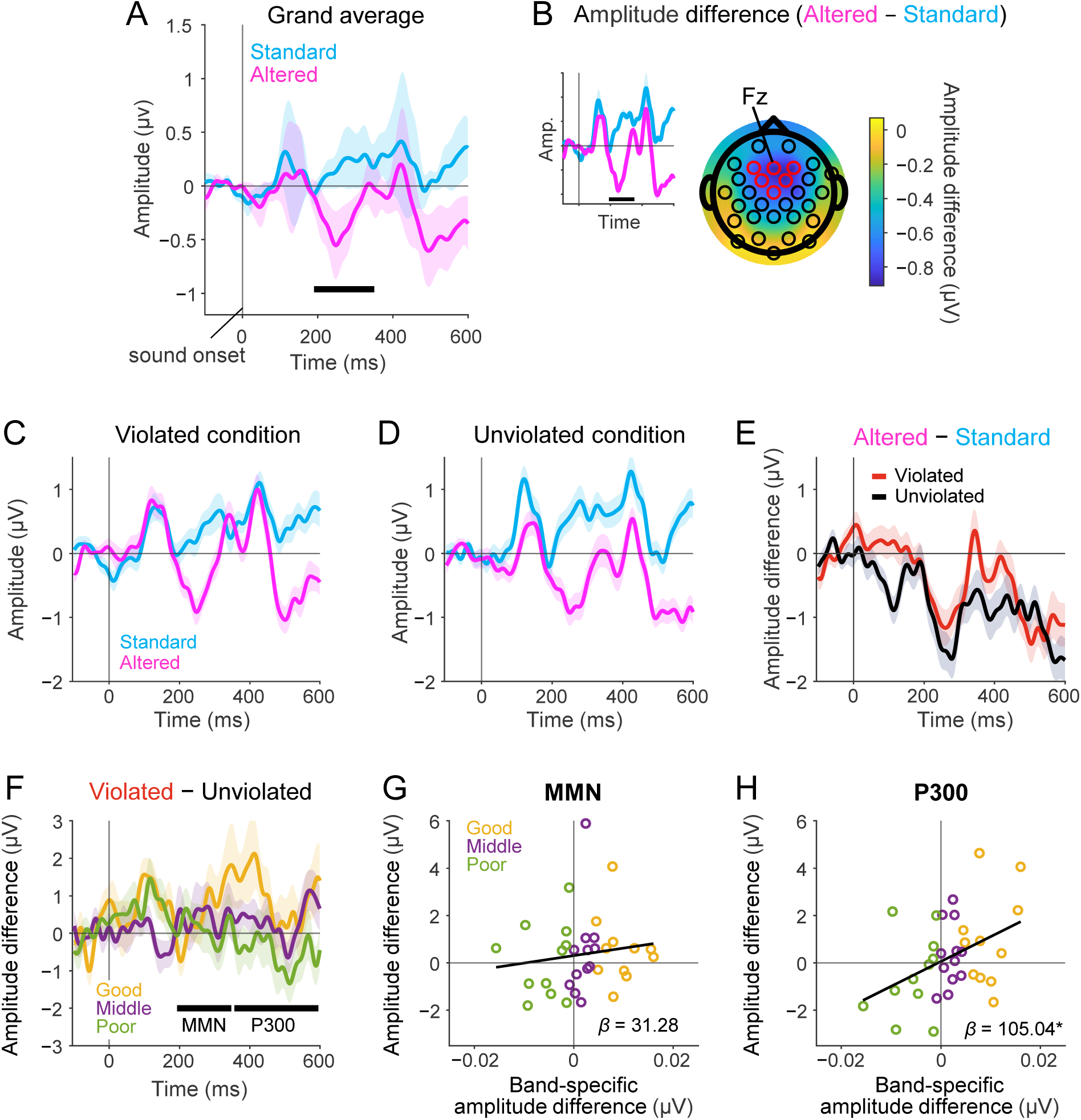
Neural responses to metrical anticipation and their relationship with entrainment. **A.** Grand average of ERPs for standard and altered sounds, without categorizing violated and unviolated conditions. The shaded areas represent SD across the electrodes. **B.** The topographical electrode map of the amplitude difference (altered − standard) shows the largest difference at the Fz electrode. Consequently, Fz and its neighboring electrodes (F3, F4, FC1, FC2, and Cz, indicated by red circles) were selected for subsequent analysis. **C,D.** ERPs in the violated (C) and unviolated (D) conditions comparing standard and altered sounds. **E.** Amplitude differences (altered − standard sounds) under both conditions. **F.** The amplitude difference (violated − unviolated) across the three participant groups determined in the analysis of the band-specific amplitude difference (see Fig. 2F). We defined the time windows at 190-350 ms and 351-600 ms after the sound onset to assess MMN and P300, respectively. The shaded area in panels C–F represents the standard error across participants. **G,H.** Scatter plots showing the relationship between the band-specific amplitude difference and the amplitude difference (violated − unviolated) for the MMN (G) and P300 (H) time windows. Colored dots represent individuals in each participant group, and the *β-*value (slope) of the regression models is indicated. Asterisk (*) indicates a significant slope (*p* < 0.05).

ERPs were then obtained by averaging the signals for both the altered and standard sounds under each condition. We subtracted the ERP elicited by standard sounds from that elicited by altered sounds to assess whether the MMN and P300 components reflected violations of the anticipatory metrical structure induced by the altered sounds, while controlling for acoustic variability. We defined the time windows at 190-350 ms and 351-600 ms after the sound onset to assess MMN and P300, respectively (shown as bold black bars in **Fig. 3F**).

#### 2.5.4. Statistics

To assess neural entrainment to meter patterns, we tested whether the amplitude differences between situations in which participants attended to triple-versus quadruple-related acoustic features were not constant across frequency bands. The data were fitted using a linear mixed-effects regression model, with amplitude difference as the dependent variable, meter-related frequency band as a fixed effect, and participant ID as a random effect as follows:

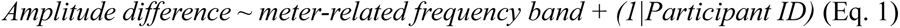

We used the ‘lmer’ function in the ‘lme4’ package (ver. 1.1.35.1) (Bates et al., 2015), and significance tests were conducted using the ‘lmerTest’ package (ver. 3.1.3) (Kuznetsova et al., 2017). After fitting the model to the data, we conducted pairwise comparisons between meter-related frequency bands using the ‘emmeans’ package (ver. 1.10.3) (Lenth, 2024). *P*-values of multiple comparisons were adjusted using the Tukey-Kramer method (denoted as *p_Tukey_*).

For the analysis of ERPs reflecting the violation of metrical anticipation, we tested whether the amplitude difference (altered versus standard sounds) in the violated condition was not the same as that in the unviolated condition. The average amplitude differences between the two conditions were compared using a paired *t*-test after confirming normal distribution. A paired Wilcoxon test was used if normality could not be confirmed.

To examine the relationship between neural entrainment and anticipatory processes, we conducted linear regression analyses. For neural entrainment, we focused on Bands-3 and 4, computing a band-specific amplitude difference by subtracting the amplitude difference in Band 4 from that in Band 3. A positive value indicates that neural activity is more strongly entrained to the triple-related feature in Band-3, whereas a negative value indicates stronger entrainment to the quadruple-related feature in the same band. For anticipatory processing, we calculated the amplitude difference between the original amplitude differences in the violated and unviolated conditions, and averaged separately for the MMN and P300 time windows. Two linear models were constructed: one with the MMN amplitude difference as the response variable, and the other with the P300 amplitude difference as the response variable, using the band-specific amplitude difference as the explanatory variable. All statistical tests were two-sided and performed using the R software (R Core Team, 2024) (version 4.1.1), with the significance level set at *α* = 0.05.

## 3. Results

### 3.1. Neural entrainment related to meter

To determine the electrodes used for the neural entrainment analysis, we computed the EEG spectra recorded during the presentation of the first 18 noise bursts, collapsing across attention manipulations (**Fig. 2A**). For each electrode, we calculated the ratio of the original to the surrogate spectral amplitude at the beat-related frequency (approximately 3.33 Hz) (**Fig. 2B**; **Table 1**). This ratio was maximal at the electrode T7. Based on this result, the signals from T7 and its neighboring electrodes (FC5, C3, and CP5) were averaged and used in all subsequent analyses.

**Table 1.** The ratio of original to surrogate EEG spectra at 3.33 Hz for each electrode.

| Electrode label | Mean ± SD |
| --- | --- |
| T7 | $-0.02 \pm 0.01$ |
| FC5 | $-0.03 \pm 0.01$ |
| C3 | $-0.03 \pm 0.02$ |
| CP5 | $-0.03 \pm 0.02$ |

Using this electrode cluster and the same time window (the first 18 noise bursts), we examined whether the participants’ EEG signals were entrained to the triple- or quadruple-related acoustic features of dual-meter stimuli. To this end, we computed the amplitude difference for each meter-related frequency band by subtracting the amplitude associated with attention to quadruple features from that associated with attention to triple features (**Fig. 2C,D**). The resulting amplitude difference values (× 10^−3^ μV, mean ± SD) for Band-1 through 5 were −1.14 ± 3.25, 0.14 ± 3.16, 2.30 ± 5.41, 0.39 ± 4.59, and −0.19 ± 3.76, respectively (**Fig. 2E**).

To assess whether these amplitude differences varied across frequency bands, we fitted a linear mixed-effects regression model (Eq.1). The model revealed a significant main effect of the meter-related frequency band (*F*(4, 132) = 3.52, *p* = 0.009). Post-hoc pairwise comparisons indicated that Band-3 exhibited significantly higher amplitude difference values than Band-1 (*p_Tukey_* = 0.004, **Table 2**). No other pairwise contrasts reached significance, although the difference between Band-3 and 5 approached significance (*p_Tukey_* = 0.073). These results indicate that, given physically identical stimuli, neural entrainment patterns shift to reflect the metrical structure that the listener perceives, with the strongest effect at the triple-meter frequency (Band-3).

**Table 2.** Pairwise comparison of overall amplitude differences between meter-related frequency bands.

| Contrast | *Estimate* ($10^{-3}$) | Lower 95% CI ($10^{-3}$) | Upper 95% CI ($10^{-3}$) | *t*-value | *p*<sub>Tukey</sub> |
| --- | --- | --- | --- | --- | --- |
| Band-1 minus Band-2 | -1.29 | -3.91 | 1.34 | -1.36 | 0.658 |
| Band-1 minus Band-3 | -3.44 | -6.07 | -0.81 | -3.62 | 0.004* |
| Band-1 minus Band-4 | -1.53 | -4.16 | 1.10 | -1.61 | 0.493 |
| Band-1 minus Band-5 | -0.95 | -3.58 | 1.67 | -1.00 | 0.853 |
| Band-2 minus Band-3 | -2.15 | -4.78 | 0.47 | -2.27 | 0.162 |
| Band-2 minus Band-4 | -0.24 | -2.87 | 2.38 | -0.26 | 1.000 |
| Band-2 minus Band-5 | 0.33 | -2.29 | 2.96 | 0.35 | 0.997 |
| Band-3 minus Band-4 | 1.91 | -0.72 | 4.54 | 2.01 | 0.266 |
| Band-3 minus Band-5 | 2.49 | -0.14 | 5.11 | 2.62 | 0.073 |
| Band-4 minus Band-5 | 0.58 | -2.05 | 3.20 | 0.61 | 0.974 |

**Note:** CI denotes confidence interval. $SE = 0.95 \times 10^{-3}$ , $df = 132$ , \* $p_{Tukey} < 0.05$

### 3.2. Event-related potentials related to anticipation

To quantify metrical anticipation, we introduced a change in either the center frequency or duration of the 19th, 20th, or 21st noise burst. These altered sounds were compared with standard sounds, defined as earlier bursts that matched the altered sounds in both center frequency and duration (**Fig. 1C**). When the participants attended to the altered acoustic feature, the modification disrupted the expected metrical structure (violated condition). When attention was directed to the other feature, the metrical structure remained intact (unviolated condition).

To select electrodes for the ERP analysis, we computed the amplitude difference between altered and standard sounds for each electrode, averaging across the conditions (**Fig. 3A,B**; **Table 3**). This difference was largest at the Fz electrode. Accordingly, data from Fz and its neighboring electrodes (F3, F4, FC1, FC2, and Cz) were averaged and used in all subsequent ERP analyses.

**Table 3.** ERP amplitude differences (altered − standard) regardless of the conditions for each electrode.

| Electrode label | Mean ± SD (μV) |
| --- | --- |
| Fz | $-0.90 \pm 0.46$ |
| F3 | $-0.76 \pm 0.36$ |
| F4 | $-0.90 \pm 0.38$ |
| FC1 | $-0.75 \pm 0.45$ |
| FC2 | $-0.89 \pm 0.46$ |
| Cz | $-0.75 \pm 0.43$ |

Employing this electrode cluster, we calculated the amplitude difference (altered minus standard) separately for the violated and unviolated conditions within the MMN and P300 time windows (**Fig. 3C–E**). In the MMN time window, the amplitude differences (μV, mean ± SD) were −0.65 ± 1.37 in the violated condition and −1.02 ± 1.51 in the unviolated condition. In the P300 time window, the corresponding values were −0.75 ± 1.39 in the violated condition and −1.03 ± 1.34 in the unviolated condition. The amplitude differences did not differ significantly between conditions for either the MMN (*V* = 351, *p* = 0.370) or the P300 (*t*(33) = 0.90, *p* = 0.374).

### 3.3. Relationship between neural entrainment and anticipatory responses

The preceding analyses separately assessed neural entrainment and anticipatory ERP responses. Next, we tested whether these two measures were related, using the band-specific amplitude difference (Band-3 minus Band-4; **Fig. 2F**) as a predictor of the amplitude difference between the violated and unviolated conditions.

To visualize individual differences in anticipatory ERP responses, we divided the participants into three equally sized groups based on their band-specific amplitude differences: good (*n* = 11), middle (*n* = 12), and poor switchers (*n* = 11) in descending order (**Fig. 2F**; no group-level statistical comparisons were performed). As shown in **Fig. 3F**, the P300 amplitude difference between the violated and unviolated conditions was graded across the three groups, with good switchers showing the largest difference. No such pattern was observed in the MMN time window.

To statistically test this relationship, we fitted linear regression models using the band-specific amplitude difference as a continuous predictor. The band-specific amplitude difference did not predict the MMN amplitude difference (*slope* = 31.28, *p* = 0.445; **Fig. 3G**, **Table 4**), but it significantly predicted the P300 amplitude difference (*slope* = 105.04, *p* = 0.013; **Fig. 3H**, **Table 5**). These results indicate that individuals with stronger meter-related neural entrainment show larger anticipatory P300 responses, suggesting a functional link between entrainment and anticipation in the perception of meter.

**Table 4.** Linear regression model to examine the impact of band-specific amplitude differences on the amplitude difference (violated minus unviolated conditions) in the MMN time window.

| Variable | *slope* [95%CI Lower, Upper] | *SE* | *t*-value | *p*-value |
| --- | --- | --- | --- | --- |
| (Intercept) | 0.31 [–0.28, 0.91] | 0.29 | 1.08 | 0.290 |
| Band-specific amplitude difference | 31.28 [–51.07, 113.63] | 40.43 | 0.77 | 0.445 |

**Note:** CI denotes confidence interval. $F(1, 32) = 0.60$ , Adjusted $R^2 = -0.01$

**Table 5.**
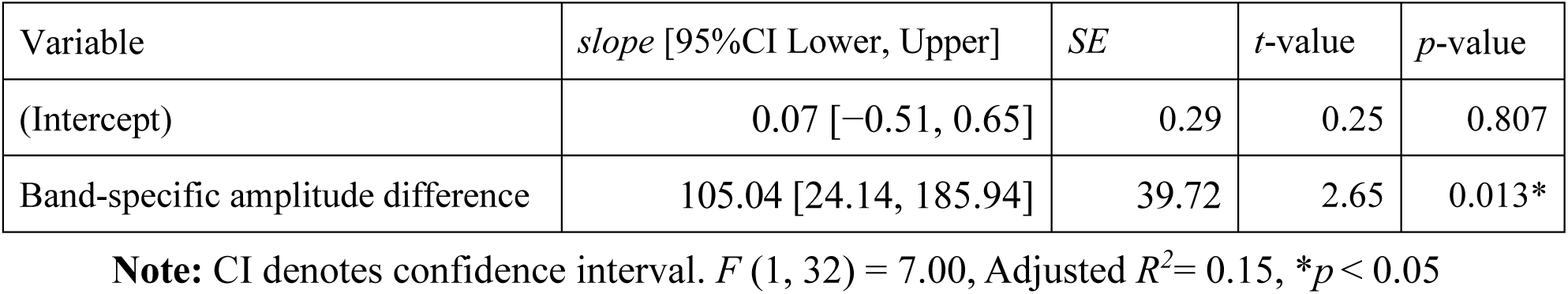
Linear regression model to examine the impact of band-specific amplitude differences on the amplitude difference (violated minus unviolated conditions) in the P300 time window.

| Variable | *slope* [95%CI Lower, Upper] | *SE* | *t*-value | *p*-value |
| --- | --- | --- | --- | --- |
| (Intercept) | 0.07 [–0.51, 0.65] | 0.29 | 0.25 | 0.807 |
| Band-specific amplitude difference | 105.04 [24.14, 185.94] | 39.72 | 2.65 | 0.013* |

**Note:** CI denotes confidence interval. $F(1, 32) = 7.00$ , Adjusted $R^2 = 0.15$ , $*p < 0.05$

## 4. Discussion

This study aimed to clarify the neural mechanisms underlying meter perception using dual-meter sound stimuli and EEG recordings. We hypothesized that selectively attending to a particular acoustic feature enhances neural entrainment to the regularity defined by that feature, and that this entrainment, in turn, gives rise to anticipation of the corresponding metrical structure. Because dual-meter stimuli support two different meters simultaneously within physically identical sound sequences, comparing neural activity across attentional conditions allows meter-related neural processes to be isolated from responses driven by acoustic input (**Fig. 1A-C**). We first examined oscillatory neural entrainment by comparing EEG spectral profiles when participants attended to either the triple- or quadruple-related features (**Fig. 2C,D**). The amplitude differences varied significantly across meter-related frequency bands, with the triple-meter band showing a larger difference than the beat-related band (**Fig. 2E**), partially consistent with our hypothesis that attention modulates neural entrainment toward the metrical structure of the attended feature. We also analyzed the MMN and P300 components of ERPs elicited by violating one’s anticipation in the metrical structure (**Fig. 3C,D**). The amplitude differences between the violated and unviolated conditions did not reach significance, and both measures showed considerable individual variability (**Fig. 3E**). This variability is informative. The band-specific amplitude difference between the triple- and quadruple-meter frequency bands (Band-3 minus Band-4), which reflects the degree to which neural entrainment matched the metrical structure of the attended feature, significantly predicted the P300 amplitude difference between conditions, but not the MMN amplitude difference (**Fig. 3F–H**), suggesting a functional link between entrainment and anticipation in meter perception. Previous studies have proposed that neural entrainment and anticipatory processing are two core mechanisms supporting meter perception (Harding et al., 2025; Snyder et al., 2024; Vuust et al., 2022) and that entrainment may serve as a basis for anticipation, as described by dynamic attending theory (Ellis & Jones, 2009; Jones, 2018; Jones & Boltz, 1989). Our study provides evidence for this proposed relationship, made possible by the combined use of dual-meter stimuli and EEG measurements.

The lack of significant effects in the band-specific amplitude differences and ERP amplitude differences between conditions is likely attributable to individual variability in top-down attentional control. In our previous study (Kondoh et al., 2021), participants practiced tapping along with each noise burst while focusing on a single acoustic feature, and those who received this training showed a greater ability to switch their perceived meter. However, because the tapping task made the metrical structure explicit, participants could have used the perceived meter to guide their attention to acoustic features, rather than deriving the perceived meter from their attentional focus on those features. To avoid the confound, the pre-training in the present study directed participants’ attention to acoustic features by having them count sounds of a given characteristic (e.g., “high” or “long”) without reference to metrical grouping. In contrast, without motor engagement, which reinforced selective attention in the previous study, individual differences in the ability to sustain attention on the instructed feature may have been amplified. In addition, the broader set of stimulus subtypes used in this study (four per category; **Fig. 1** and **Supplementary Fig. S1**) may have made it more difficult for some listeners to maintain their attention consistently across trials. Musical experience may also have contributed to the observed variability (Drake et al., 2000; Geiser et al., 2009; Kondoh et al., 2021), although we did not assess it using standardized measures, such as the Goldsmiths Musical Sophistication Index (Müllensiefen et al., 2014; Sadakata et al., 2022). Future research should systematically evaluate these factors to clarify the sources of individual differences in neural entrainment and anticipatory responses.

The band-specific amplitude differences reflecting neural entrainment did not predict the amplitude differences between the violated and unviolated conditions in the MMN time window (**Fig. 3G**), but significantly predicted those in the P300 time window (**Fig. 3H**). This dissociation is consistent with the functional distinction between the two components. MMN is typically associated with the pre-attentive detection of acoustic deviations (Koelsch et al., 2019; Näätänen et al., 1978, 2007; Vuust et al., 2022), and previous studies have shown that disruptions to low-level acoustic regularities, such as amplitude patterns in a duple meter, can elicit MMN responses (Radchenko et al., 2023; Zhao et al., 2017). In the present study, the metrical violations involved an abstract grouping structure that depends on sustained selective attention, which may not be readily captured by the pre-attentive processes indexed by MMN. In contrast, the P300 component reflects the conscious evaluation of unexpected events (Koelsch et al., 2019; Polich, 2007; Vuust et al., 2009). Vuust et al. (2009) showed that violations of rhythmic expectations evoke stronger P300 responses in jazz musicians than in non-musicians, indicating that P300 amplitude is sensitive not merely to acoustic deviance but to higher-order metrical expectations shaped by experience. Our finding that P300 responses were predicted by the strength of neural entrainment is consistent with this view, and further suggests that such expectations can arise from oscillatory synchronization with the metrical regularity from the attended acoustic feature.

Previous studies on neural entrainment related to meter perception have commonly relied on steady-state evoked potentials (SSEPs), in which the amplitude of the EEG signal at the frequency corresponding to the perceived meter is isolated by subtracting the mean amplitude of the neighboring frequency bins (Chemin et al., 2014; Cirelli et al., 2016; Gibbings et al., 2023; Nave et al., 2022; Nozaradan et al., 2011, 2012, 2016, 2018). While effective for identifying frequency-specific neural activity, the SSEP approach normalizes each frequency bin against its local spectral neighborhood, which can make it difficult to directly compare the relative strength of entrainment across different meter-related frequencies. The present study addressed this limitation by comparing EEG spectra between attentional conditions in which the participants listened to physically identical stimuli. By subtracting one condition from the other, the acoustic contributions are canceled, enabling a direct comparison of the entrainment strength across meter-related frequency bands on a common scale. Therefore, this spectral subtraction method may complement traditional SSEP techniques in studying attention-based neural entrainment to meters.

The present study had several limitations. First, we used dual-meter stimuli with IOIs fixed at 300 ms (200 beats per minute or bpm) to maintain consistency with our previous study (Kondoh et al., 2021). This setting is unlikely to be problematic for beat perception because earlier research has indicated an optimal tempo for beat perception ranging from 100 ms to 2 s for the IOI (London, 2012; Repp & Su, 2013). However, 200 bpm could be considered very fast in Western musical standards; thus, further research employing slower tempos is necessary to clarify its effects on neural entrainment and anticipatory responses during meter perception. Second, we did not rigorously assess the relationship between band-specific amplitude differences (reflecting neural entrainment related to meters), amplitude differences between conditions (reflecting expectancy violations), and behavioral measures of meter perception. As an exploratory analysis, we examined the correlations between task sensitivity, measured by the ability to detect whether an altered sound disrupted metrical structures, and EEG indicators (**Supplementary Text S1, Fig. S3**). However, because we did not have a priori hypotheses about specific electrode sites, we only reported electrodes with high correlation coefficients. Future studies should systematically test the relationship between EEG and behavioral indicators using the electrodes identified in our supplementary analyses. Third, the regression analysis revealed a significant association between the entrainment index and the P300 amplitude difference, with an observed effect size in the medium range (f^2^ = 0.22, calculated as *F* × *df_1_*/*df_2_* = 7.00 × 1/32). This value fell slightly below the minimum detectable effect size identified by the sensitivity analysis (f^2^ = 0.25 at 80% power), suggesting that the present sample was near the boundary required to detect this effect at conventional power levels. Replication with larger samples would strengthen confidence in the reliability of this association.

In conclusion, this study investigated the neural basis of meter perception using dual-meter stimuli designed to minimize the confounding effects of acoustic variations on neural activity. We hypothesized that top-down selective attention to either the frequency or duration of a sound sequence modulates neural synchronization with the acoustic regularities of that feature, giving rise to expectations that support meter perception. Consistent with this hypothesis, the strength of meter-related neural entrainment predicted the magnitude of anticipatory P300 responses to such violations. These findings represent a methodological advance in dissociating neural responses to metrical structure from acoustic confounds, and provide evidence linking neural entrainment to anticipatory processing, a fundamental question in human auditory cognition.

## CRediT authorship contribution statement

**Sotaro Kondoh:** Conceptualization, Data Curation, Formal Analysis, Funding Acquisition, Investigation, Methodology, Software, Visualization, Writing – Original Draft Preparation, Writing – Review & Editing.

**Kazuo Okanoya:** Conceptualization, Funding Acquisition, Project Administration, Supervision, Writing – Review & Editing.

**Ryosuke O. Tachibana**: Conceptualization, Formal analysis, Funding Acquisition, Methodology, Project Administration, Software, Supervision, Visualization, Writing – Review & Editing.

## Data accessibility

The data supporting the findings of this study as well as the sound files of the dual-meter stimuli used are available from the Open Science Framework repository (https://osf.io/hw2y8).

## Declaration of Competing Interest

The authors have declared that no competing interests exist.

## Acknowledgments

This work was supported by grants from JSPS KAKENHI (grant numbers 23H05428 to KO, 23K18475 and 24H00735 to ROT, and 24KJ1930 to SK). The authors would like to thank Drs. Hironori Nakatani and Reiko Shiba for their valuable feedback on the early versions of our pilot experiment. We would also like to thank Dr. Shinya Fujii and the members of his laboratory for their insightful comments on the preliminary results of this study.

## Scientific transparency statement

**DATA**: All raw data supporting this research are publicly available: https://osf.io/hw2y8. Processed data will be publicly available before acceptance.

**CODE:** All analysis code supporting this research will be publicly available before acceptance: https://osf.io/hw2y8.

**MATERIALS:** All study materials supporting this research are publicly available: https://osf.io/hw2y8.

**DESIGN:** This article reports, for all studies, how the author (s) determined all sample sizes, all data exclusions, all data inclusion and exclusion criteria, and whether inclusion and exclusion criteria were established prior to data analysis.

**PRE-REGISTRATION:** No part of the study procedures was pre-registered in a time-stamped, institutional registry prior to the research being conducted. No part of the analysis plans was pre-registered in a time-stamped, institutional registry prior to the research being conducted.

## Supplemental Materials

### Supplementary Text S1. Exploring the relationship between neural entrainment, anticipation, and behavioral sensitivity in detecting metrical violations

To analyze the relationship between neural indices (entrainment and anticipation) and behavioral sensitivity to violations of metrical expectations, we conducted a supplemental behavioral analysis. After the EEG session, participants completed a task in which they judged whether an altered sound introduced a deviation when attending to either its center frequency or its duration. Each participant evaluated 28 stimuli per acoustic feature, resulting in 56 randomized trials. Participants had 2.5 s to respond and were instructed to answer as quickly and accurately as possible. If no responses were recorded, the next trial began automatically.

Sensitivity to metrical violations was quantified using the signal-detection theory (Gescheider, 1997). A hit was defined as the correct detection of a violation, whereas a miss indicated a failure to detect it. A false alarm referred to an incorrect report of a violation when none was present, and a correct rejection was an accurate identification of no violation. We define *Z*_*N*_ as the z-transformed value of (1 minus the proportion of false alarms), and *Z*_*SN*_ as the z-transformed value of (1 minus the proportion of hits). Detection accuracy (*d*′) was calculated using the following equation:

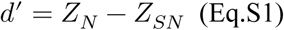

We then examined the relationship between *d*′ (*mean* = 0.29, *SD* = 0.47) and three neural measures: (i) band-specific amplitude differences, (ii) amplitude differences (between the violated and unviolated conditions) during the mismatch-negativity (MMN) time window, and (iii) amplitude differences in the P300 time window. Because we had no predefined hypotheses about electrode locations, we computed the Pearson correlation coefficients without formal significance testing.

Stronger positive correlations between *d*′ and band-specific amplitude differences were observed at F7 (*r* = 0.38), FC6 (*r* = 0.32), and FC5 (*r* = 0.29) (**Supplementary Fig. S3A**). During the MMN window, the left hemisphere electrodes showed relatively strong negative correlations between *d*′ and the amplitude difference at CP5 (*r* = −0.30), P3 (*r* = −0.29), and FC1 (*r* = −0.25) (**Supplementary Fig. S3B**). In the P300 window, stronger positive correlations were observed in the right hemisphere electrodes such as FT10 (*r* = 0.40), PO10 (*r* = 0.35), and T8 (*r* = 0.34) (**Supplementary Fig. S3C**). Therefore, these electrode sites may serve as promising targets for future studies investigating how neural indices of entrainment and anticipation relate to behavioral indices of meter violation detection.

---

**Supplementary Fig. S1.**
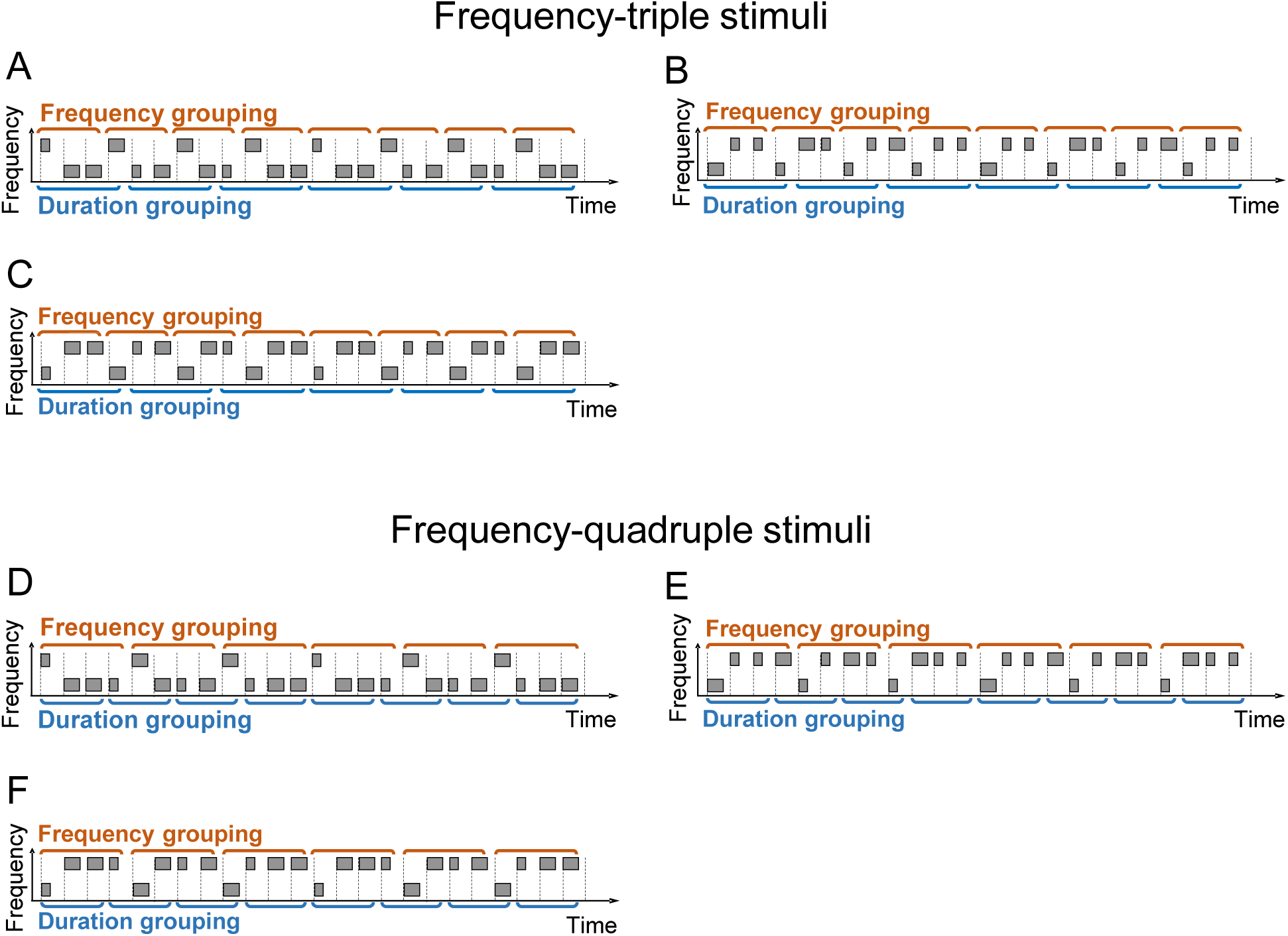
Schematic representation of dual-meter stimuli not included in the main text. Each gray square represents a band-limited noise burst, with each stimulus consisting of 24 bursts. The center frequency of the higher sound was 1600 Hz and that of the lower sound was 1400 Hz, resulting in a frequency difference of 200 Hz. The duration of the longer sound was 150 ms, whereas that of the shorter sound was 50 ms, creating a duration difference of 100 ms. The bandwidth of each noise burst was set to 120% of its center frequency, and the inter-onset intervals (IOIs) were fixed at 300 ms. **A-C.** Frequency-triple stimuli. The frequency pattern is either “high–low–low–…” (panel A) or “low–high–high–…” (panels B and C), while the duration pattern follows either “short–long–long–long–…” (panels A and C) or “long–short–short–short–…” (panel B). **D-F.** Frequency-quadruple stimuli. The frequency pattern follows “high–low–low–low–…” (panel D) or “low–high–high–high–…” (panels E and F), while the duration pattern is either “short–long–long–…” (panels D and F) or “long–short–short–…” (panel E).

**Supplementary Fig. S2.**
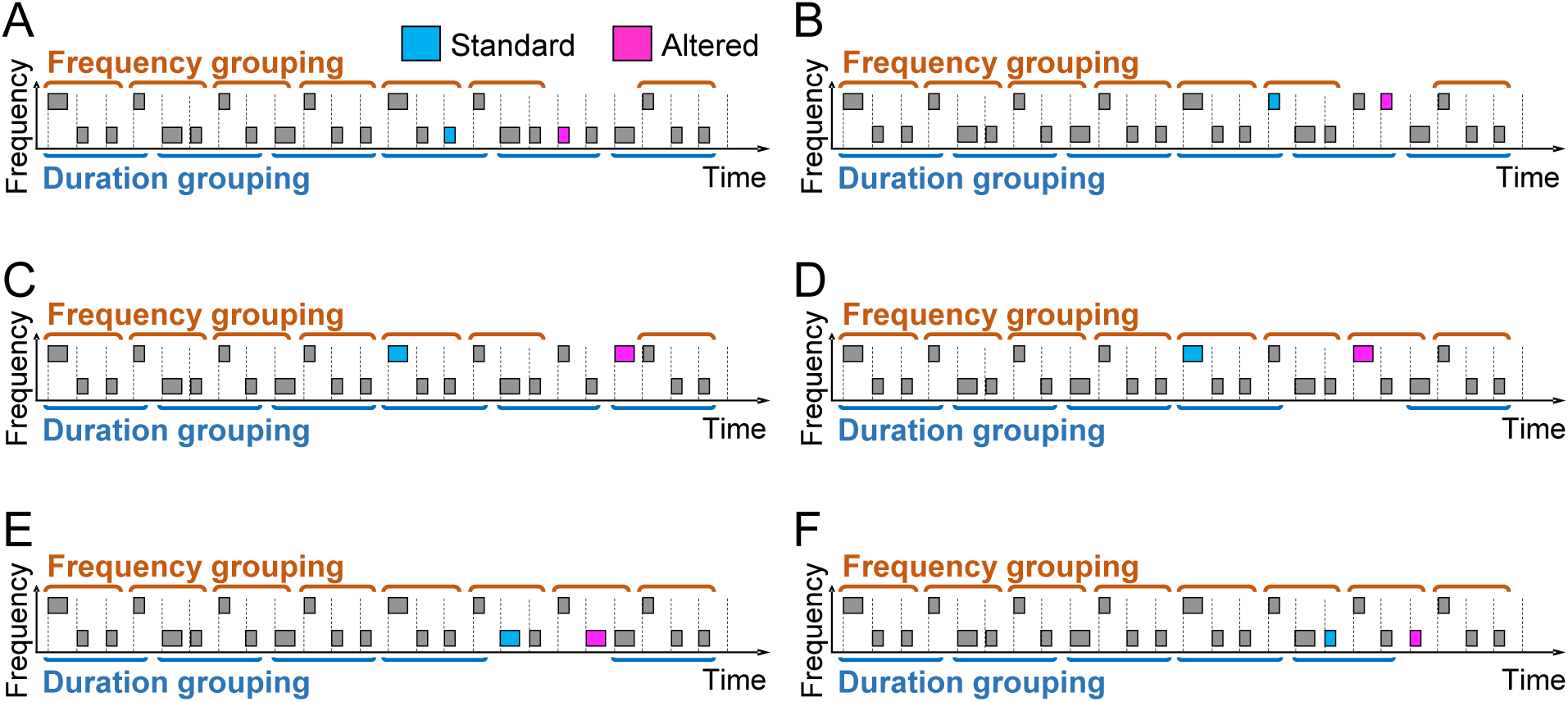
Variations in one stimulus (Fig. 1A in the main text). Each pink square represents an altered sound, either by changing the center frequency of the 19th (panel A: high to low), 20th (panel B: low to high), or 21st (panel C: low to high) noise bursts or by changing the duration of the 19th (panel D: short to long), 20th (panel E: short to long), or 21st (panel F: long to short) noise bursts. For EEG analysis, standard sounds were defined as those that matched the altered sounds in both center frequency and duration and were presented at least three sounds ahead (indicated by light blue squares).

**Supplementary Fig. S3.**
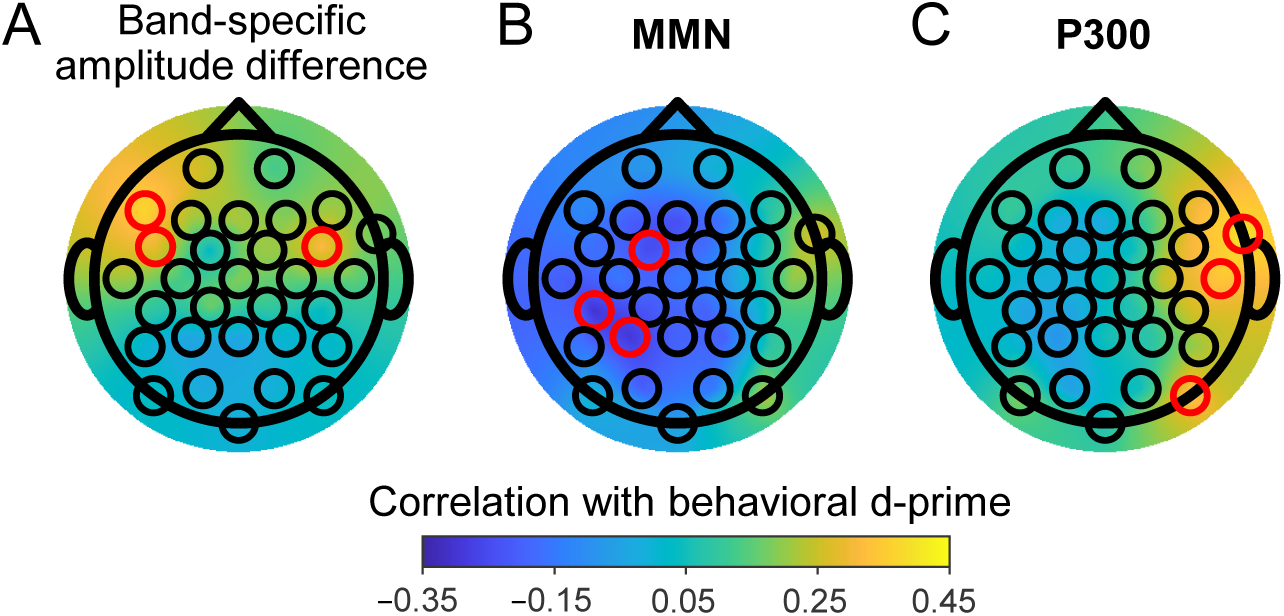
Topographical distribution of correlations between behavioral d-prime (*d*′) and EEG indices across the electrodes. **A.** Band-specific amplitude difference. The three electrodes (F7, FC5, and FC6, indicated by red circles) showed stronger positive correlations. **B.** Amplitude difference in the MMN time window. The three electrodes (CP5, P3, and FC1, indicated by red circles) exhibited stronger negative correlations. **C.** Amplitude difference in the P300 time window. The three electrodes (FT10, PO10, and T8, indicated by red circles) showed stronger positive correlations.

## Notes

https://osf.io/hw2y8

### Competing Interest Statement

The authors have declared no competing interest.

### Summary of Updates

The Introduction and Discussion were revised to improve logical flow. In the Results, descriptive statistics were relocated to tables to enhance the readability of key inferential statistics.

